# HCAtk and pyHCA: A Toolkit and Python API for the Hydrophobic Cluster Analysis of Protein Sequences

**DOI:** 10.1101/249995

**Authors:** Tristan Bitard-Feildel, Isabelle Callebaut

## Abstract

Motivation: Detecting protein domains sharing no similarity to known domains, as stored in domain databases, is a challenging problem, particularly for unannotated proteomes, domains emerged recently, fast diverging proteins or domains with intrinsically disordered regions.

Results: We developed pyHCA and HCAtk, a python API and standalone tool gathering together improved versions of previously developed methodologies, with new functionalities. The developed tools can be either used from command line or from a python API.

Availability: HCAtk and pyHCA are available at https://github.com/T-B-F/pyHCA under the CeCILL-C license.

## Introduction

The annotation of a protein sequence is very often the first step of many bioinformatics analyses, for instance for studying the function of a gene or the evolution of organisms. Protein domain annotation dominates analyses, describing a protein as a list of blocks corresponding to evolutionary and functional conserved segments. Protein domain families have been extensively compiled through sequence or structure similarity searches and stored in several public databases. These domain databases represent state of the art of our current knowledge of the protein domain universe [11, 18]. However, many protein sequences escape, at least partially, domain annotation, particularly in non-model organisms and remain in the so-called dark protein sequence universe [4]. Classical methodological approaches model protein domain families as Hidden Markov Models (HMMs). However, to that aim, sequences need to be clustered and aligned based on the identification of sequence similarities. Therefore, proteins from organisms distant from the species considered in the model, fast diverging proteins, recently emerged domains and domains containing disordered regions, are less likely to be annotated using this methodology [5]. Here, in order to provide a comprehensive view of protein domain architectures, we present a standalone software named HCA toolkit (HCAtk), and its associated python API pyHCA. The package is easily installable and extend our previous developed tools making use of the Hydrophobic Cluster Analysis (HCA) of protein sequences [6, 8–10, 12, 24], with new functionalities. The HCA methodology, based on a two-dimensional representation of protein sequences, highlights clusters of hydrophobic amino acids making up globular domains. More on the HCA methodology can be found in the supplementary materials.

## Methods

Seg-HCA [10] was developed to automatically delineate potential “foldable” domains within protein sequences and is the core part of our package. Recently, Piovesan et al. [21] implemented an in-house version of Seg-HCA in FELLS, which allows to nicely visualize different properties of a protein sequence. Our new version of Seg-HCA was rewritten for speed and a score is now computed, describing the general composition in hydrophobic clusters of the delineated foldable domains. This score is compared to an empirical distribution computed over 734 disordered protein sequences from DisProt v7 [20] to produce a p-value. Figure 1A shows the distributions of scores computed using non redundant sequences of the Protein Data Bank for the set of globular domains and the set of DisProt protein sequences. The resulting p-value can thus be used to evaluate the likelihood of the delineated domains to fold into globular structures. Interestingly, some Seg-HCA domains are reported with a high p-value. A closer inspection revealed these sequences as partially disordered and undergoing possible folding upon binding. A detailed description of scores with some examples is provided in the supplementary material.

**Figure 1:**
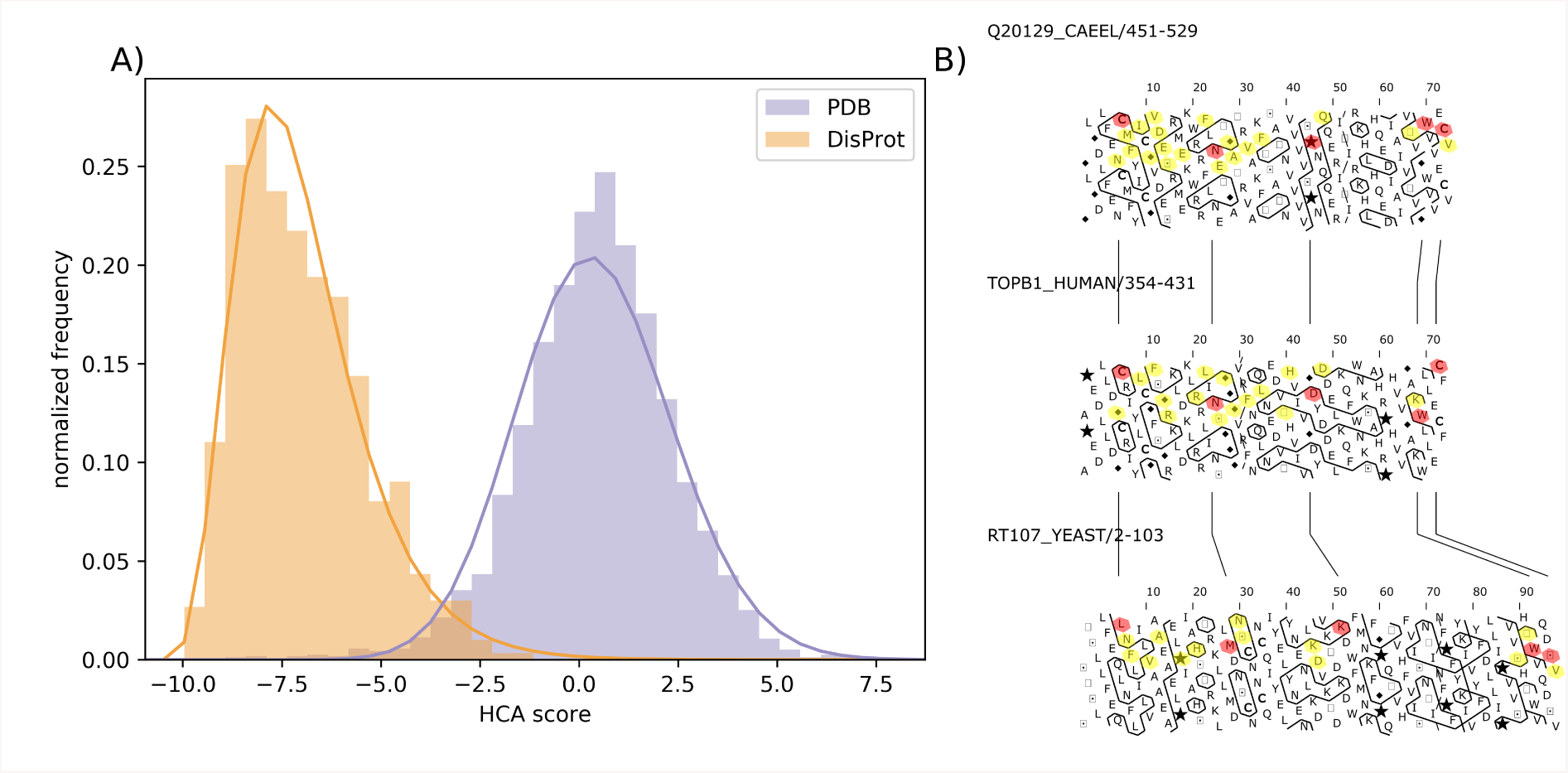
HCA score and HCA plot example. Panel A, left, shows the normalized HCA score distribution calculated for protein sequences from DisProt v7 (left, orange { disordered sequences) and from PDB (right, violet - globular domains). The HCA p-value assessing the globularity of delineated foldable segment, is computed using the empirical distribution from DisProt sequences. Panel B, right, shows the HCA plots of three BRCT domains from the Pfam family (PF00533). The aligned protein sequences were used as an input and conserved amino acids can be visualized in red (highly conserved) and yellow, in the context of hydrophobic clusters (HC), in order to evaluate the secondary structure conservation, relatively to the HC shapes.

The second methodology included in the package is our TREMOLO-HCA software (Traveling through REMOte homoLOgy) [9]. Using as queries domains delineated using Seg-HCA, remote similarity search is performed against protein sequences from the Uniprot database [29] using HHBlits [25]. For each hit, domain arrangement of the Uniprot targets is retrieved from the Interpro database [17]. The final output allows to directly link unknown protein domains, delineated by Seg-HCA, to existing annotations and to analyze these unknown domains in the context of their domain arrangement. The original tool was based on PSI-Blast and the CDD webserver. The new implementations based on HHBlits and Interpro allows a more sensitive detection of protein sequence remote similarity combined with a larger coverage of the protein domain universe thanks to the multiple sources of annotations integrated in Interpro. Several scripts are also provided to easily parse and query the TREMOLO output and to quickly retrieve protein domains of Interpro overlapping the unknown Seg-HCA domains or to retrieve the whole protein domain architectures associated with the Seg-HCA domains.

Finally, two drawing functionalities were developed. The first functionality allows visualization of hydrophobic clusters of protein sequences, whether these are aligned or not. For each protein provided in a fasta file, an HCA plot is drawn, allowing the quick inspection of the hydrophobic cluster content of a protein sequence, which gives information about its composition in regular secondary structures. A detailed description of the drawing methodology is provided in the Supplementary Material. Moreover, another new functionality was implemented to highlight conservation between aligned protein sequences on their HCA plots. Conserved protein sequence positions can therefore be inspected in the context of their hydrophobic cluster organization (Figure 1B). The second drawing functionality is a new methodology built on the TREMOLO results to easily visualize the known protein domain annotation (from Interpro) and the newly delineated domains in an evolutionary context by using the NCBI taxonomic database. The tree is automatically built by fetching the taxonomic id of the Uniprot target sequences found by TREMOLO thanks to the ete3 python package. The Seg-HCA domains of TREMOLO can then be analyzed in the context of their protein domain arrangement and visualized in terms of taxonomic specificity and domain association (Fig. S1). The HCA toolkit is written in python3 and is provided under the CeCILL-C license agreement. The functions associated with the HCA analyses in the toolkit can also be directly used through a python API and as such can easily be used in other software.

## Supporting Information

Supplementary Figure 1 is accessible at https://github.com/T-B-F/pyHCA/blob/master/img/Supplementary_Fig1.pdf.

### HCA methodology, HCA plot and Seg-HCA

HCA hydrophobic clusters, made of strong hydrophobic amino acids (V, I, L, F, M, Y, W), are different from hydrophobic segments as they can incorporate other, non-hydrophobic residues. This property originates from the use of a two-dimensional alpha-helical net, connecting hydrophobic amino acids separated by up to three non-hydrophobic amino acids (or a proline) [12]. Hydrophobic clusters defined in this way (with this hydrophobic alphabet and the connectivity distance associated with the *α*-helix) have been shown to match at best regular secondary structures (*α*-helices and *β*-strands) and to constitute hallmarks of folded domains [8, 30]. Sequence segments delineated by Seg-HCA, which correspond to regions where a high density in hydrophobic clusters is detected, have been shown to correspond to domains that have the ability to fold, either in an autonomous way or following contact with partners [5, 10]; these segments are later referred to as HCA domains. The advantage of Seg-HCA for the characterization of the dark proteome is to allow the prediction of these foldable domains from the only information of a single amino acid sequence, without the prior knowledge of homologous sequences.

Figure S2 presents the Hydrophobic Cluster Analysis (HCA) methodology and indicates how are generated the HCA plots shown in Figure 1B. From an original 1D amino acid sequence (panel A), a 2D plot is created (panel D) by rolling the amino acid sequence around an *α*-helix (panel B) and cutting the helix along the horizontal axis. The helix forms a plane (panel C) on which every line of amino acids corresponds to an helix turn. The plane is duplicated and the hydrophobic clusters are defined by joining contiguous strong hydrophobic amino acids (V, I, L, F, M, Y, W).

Regular secondary structure (RSS) elements can be easily visualized on the 2D plot, mainly corresponding to hydrophobic clusters, which are separated from each other by loops. Vertical hydrophobic clusters mainly correspond to *β*-strands and horizontal clusters to *α*-helices. A dictionary of the most current hydrophobic clusters, established from a comprehensive analysis of experimental 3D structures, can be considered for evaluating the main propensities of hydrophobic clusters towards RSS [8, 24].

### HCA score

The HCA score, used to compute a p-value associated with each HCA domain, is defined as follow. Each residue of an HCA segment is associated with a class regarding the residue type and hydrophobicity. Such a residue is either in an hydrophobic cluster and hydrophobic, in an hydrophobic cluster and hydrophilic, or outside an hydrophobic cluster. A value is attributed to each class and the HCA score is computed as follow: with s(i) the function mapping the residue i to each class value.

Therefore, the HCA score scales with the density in hydrophobic clusters and in hydrophobic residues inside the clusters. As the HCA score calculation motivation is to provide an estimation of the globular character, i.e. the foldability of an HCA domain, the value of each of the three classes was optimized to maximize the separation between the distributions of the HCA scores computed on disordered sequences

**Figure S2:**
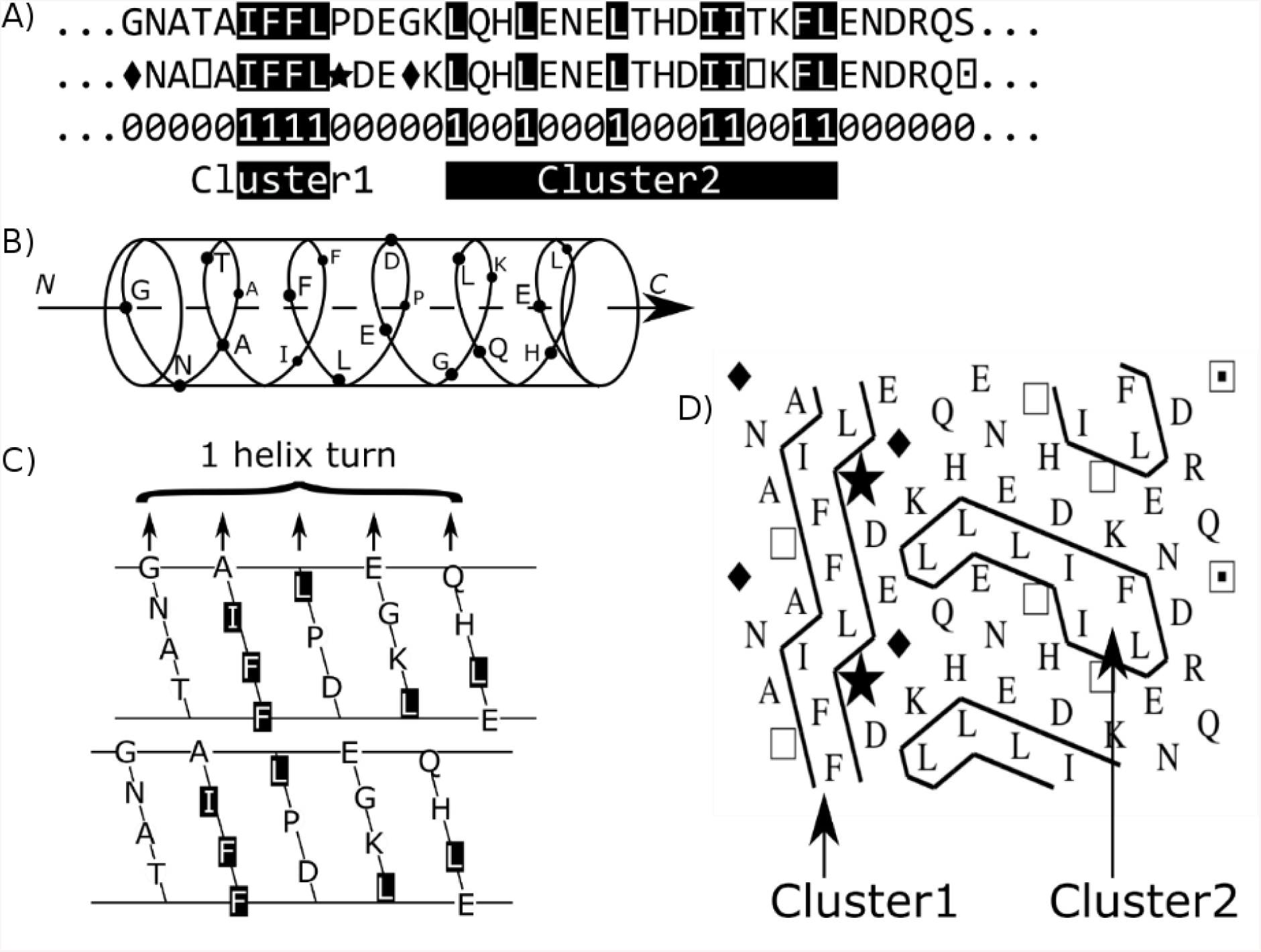
Principle of the HCA plot. Panel A, the protein sequence (1D), in which hydrophobic amino acids are represented as white letters, is written on an *α*-helix, displayed on a cylinder (panel B). This one is cut along the horizontal axis and unrolled, in order to get the full environment of each amino acid, as it exists on the 1D sequence (panel C). Strong hydrophobic amino acids (V, I, L, F, M, Y, W) are encircled and their contours are joined (panel D), forming clusters. Horizontal and vertical clusters are mainly associated with alpha helices and beta strands, respectively.

 from DisProt and on sequences of globular proteins from the PDB. The optimization was performed on a grid search of every combination of integers between [-10; 10]. The best combination was obtained with the values of 10, 9 and -10 for classes of residues in an hydrophobic cluster and hydrophobic, in an hydrophobic cluster and hydrophilic and outside an hydrophobic cluster, respectively.

Finally, a reciprocal inverse Gaussian random distribution was fitted on the DisProt scores distribution and the cumulative density function of this random distribution was used to define the p-value associated to each HCA domain. To avoid problems of scores associated with short HCA domains (≤ 30 residues), and as the limited number of sequences in DisProt makes difficult to adapt the scores to various sequence lengths, the p-values are not reported in these particular cases. This methodological choice is justified as a minimal number of residues is generally necessary for a protein domain to fold into a globular structure.

### Examples of disordered regions with low HCA scores

Supplementary Figures 3 and 4 show two examples of protein sequences taken from the left tail of the HCA score distribution of DisProt sequences, displayed in Figure 1A of the main document. For each figure, the HCA plot is drawn on top and the DisProt annotation taken from the DisProt webserver is shown at the bottom. These two sequences have HCA patterns typical of disordered proteins, i.e. proteins having very few hydrophobic residues, often gathered in HCA clusters of small length and spread along the sequence, this one including many proline residues (star symbols). Both proteins regions shown in Fig. S3 and S4 have low percentages of hydrophobic amino acids (13% and 6%).

**Figure S3:**
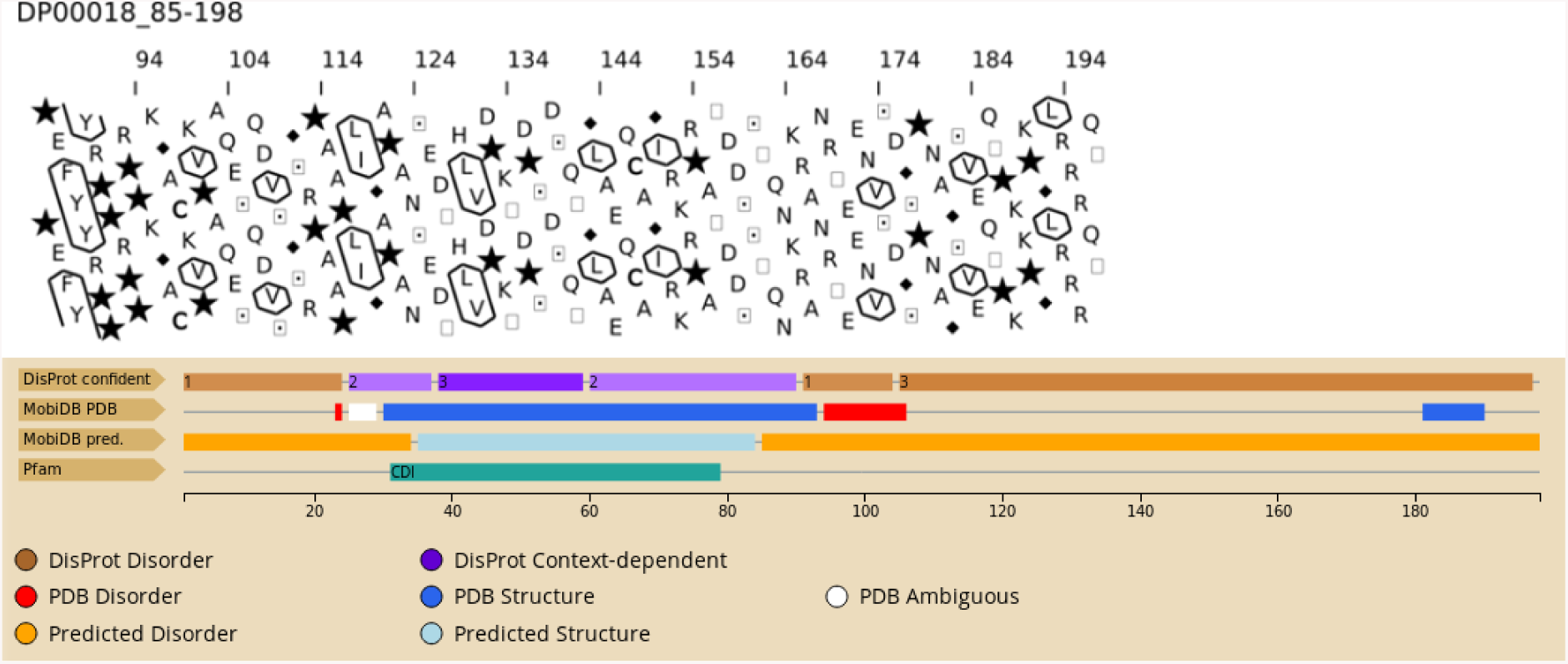
Disordered regions with low HCA score - example n° 1. The HCA pattern displayed by the disordered region is typical of non globular regions, with very few clusters and many proline (star symbols). The disordered region corresponds to the sequence segment from amino acids 85 to 198 of the human Cyclin-dependent kinase inhibitor p27(Kip1) (Uniprot P46527).

Figure S3 corresponds to the C-terminal sequence of the human Cyclin-dependent kinase inhibitor p27. p27Kip1 controls eukaryotic cell division through interactions with cyclin-dependent kinases [22] and is known as a flexible protein [13], whose stability is associated with phosphorylation. The C-terminal region of p27 has a high flexibility, which provides the molecular basis for the sequential signal transduction conduit that regulates its own degradation and cell division [3, 13]).

Figure S4 corresponds to the C-terminal sequence of the chicken Histone H5 protein. Histone proteins have well characterized intrinsic disordered regions which are necessary to their biological function [19] and are targets for post-translational modifications recognized by specific readers. Two serine phosphorylation sites have been identified at position 146 and 167. The abundance of lysine and arginine also suggests possible acetylation/methylation sites. On the other hand, the C-terminal domain of chicken Histone H5 has a DNA binding motif [23], whose activity requires a high level of intrinsic flexibility.

**Figure S4:**
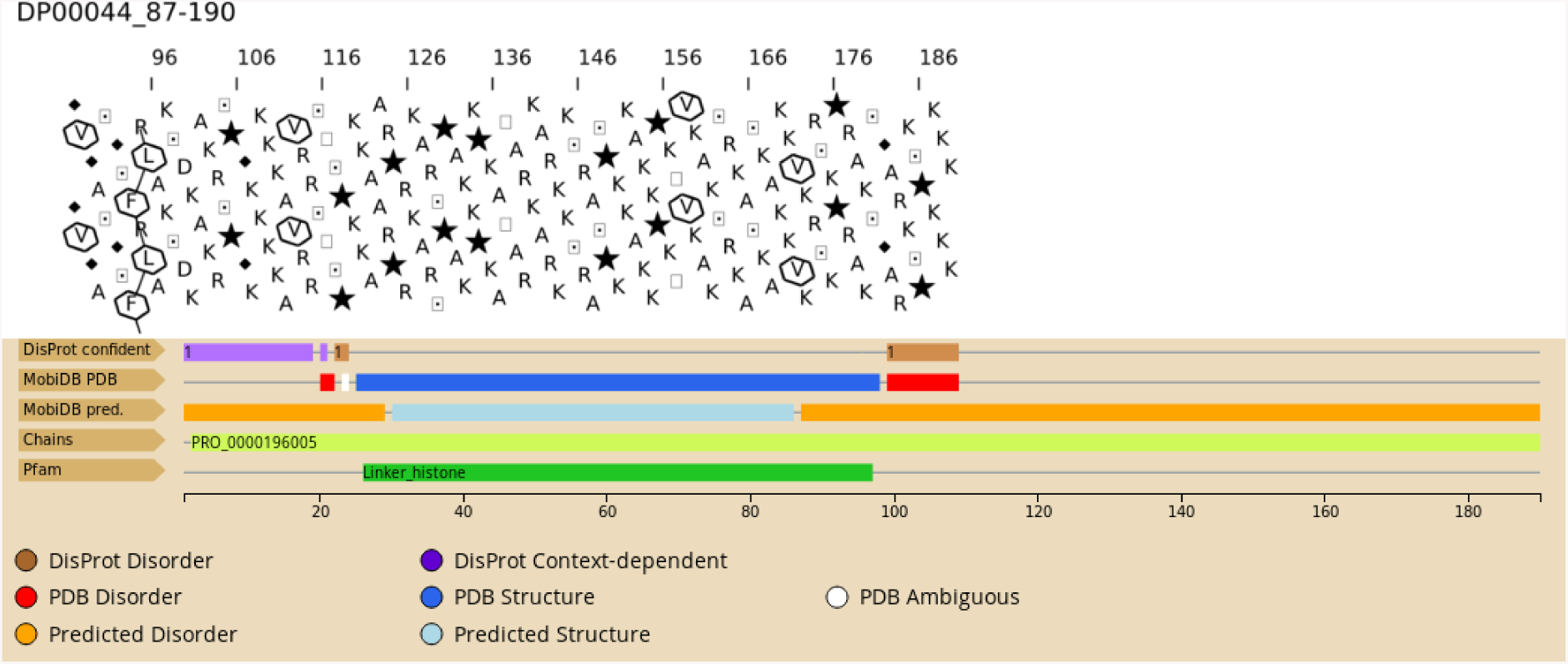
Disordered regions with low HCA score - example n° 2. The HCA pattern displayed by the disordered region is typical of non globular regions, with very few clusters and many proline (star symbols). The disordered region corresponds to the sequence segment from amino acids 87 to 190 of the chicken Histone H5 protein (Uniprot P02259).

### Examples of disordered regions with high HCA scores

Figures S5 and S6 show two examples extracted from the right tail of the DisProt HCA score distribution, i.e. proteins with HCA scores similar to the lowest scores of the sequences extracted from PDB (named PDB sequences below). These two examples display HCA patterns including larger hydrophobic clusters (typical of regular secondary structures), as found in globular proteins, but with a slightly lower total content in hydrophobic amino acids (24% for both against 30% expected). The first example concerns a disordered region (amino acids 291 to 352) found in the chicken zing finger FYVE domain-containing protein 9 (UniProt O95405, Fig. S5). The 3D structure of only the FYVE domain has been experimentally characterized, corresponding to the FYVE zinc finger domain (PF01363) (amino acids 663 to 751), the second domain corresponds, from amino acids 1048 to 1400, to a Pfam domain of unknown function (PF11979). The protein regulates the subcellular localization of SMAD2/SMAD3 by recruiting them to the TGF-beta receptor [7, 28]. The HCA pattern displayed by the disordered region is similar to patterns observed for foldable regions, suggesting that this small domain is able to fold, at least under particular conditions. The absence of any clear annotation in the N-terminal part of the protein, including two small regions predicted as disordered, but in which a potential order is detected, suggests the presence of an un-detected domain of unknown function [4]).

**Figure S5:**
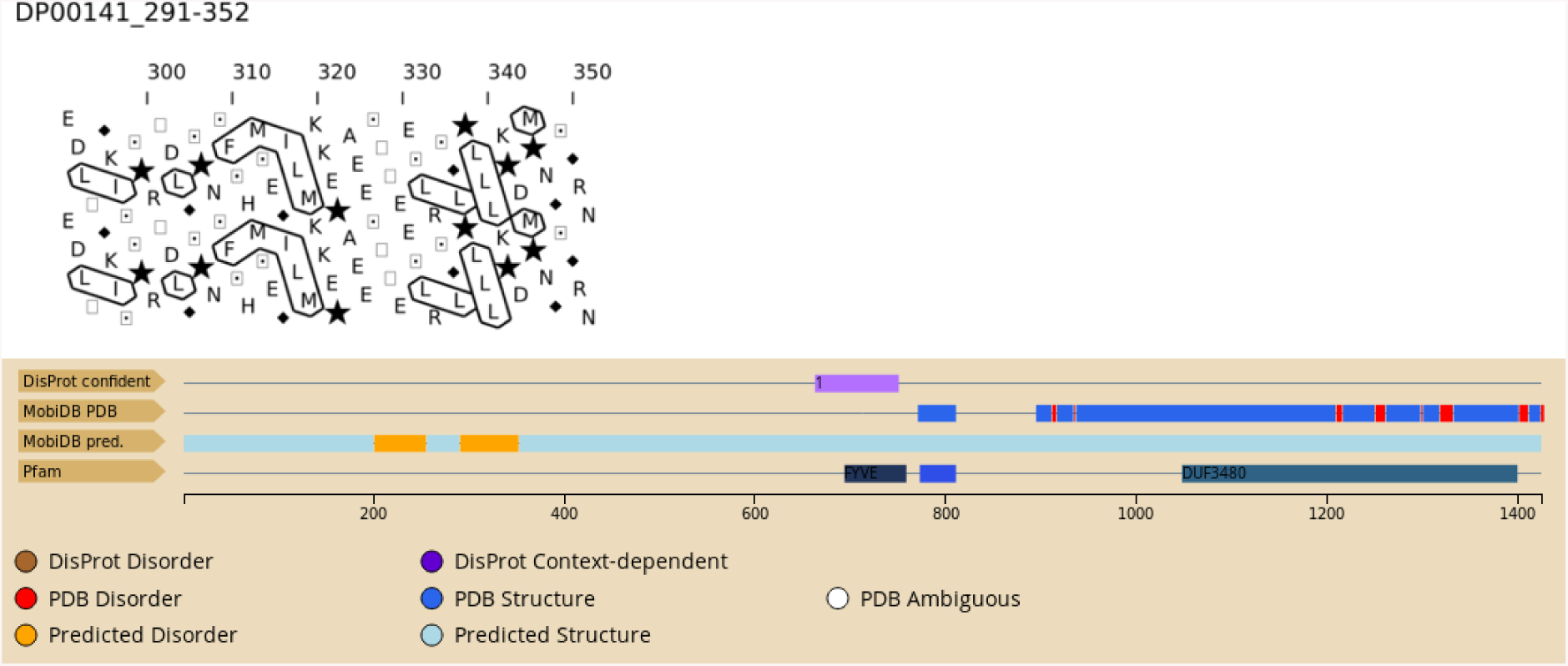
Disordered regions with high HCA score - example n°1. The HCA pattern displayed by this disordered region is similar to patterns observed for foldable regions. Clusters of hydrophobic residues, whose length are typical of stable regular secondary structures, are separated by regions whose lengths are typical of loops. This disordered region corresponds to an internal segment, from amino acids 291 to 352, of the chicken’s zinc finger FYVE domain-containing protein 9 (UniProt O95405), without known function.

The second example concerns a disordered region (amino acids 82 to 134) found in another member of the zinc finger family (zinc finger protein ZNF593, UniProt O00488, Fig. S6) which belongs to the human species. The zinc finger domain is central (60 to 86) and flanked by two disordered regions. The protein has a high degree of intrinsic disorder, as revealed by the experimental NMR structure of the full length protein, without truncation of the N- or C-terminal regions [15]. No clear function is currently associated with the N- and C-terminal regions of the protein. As for the previous example, the HCA pattern displayed by the disordered region is similar to patterns observed for foldable regions, suggesting that this small domain is able to fold, at least under particular conditions.

**Figure S6:**
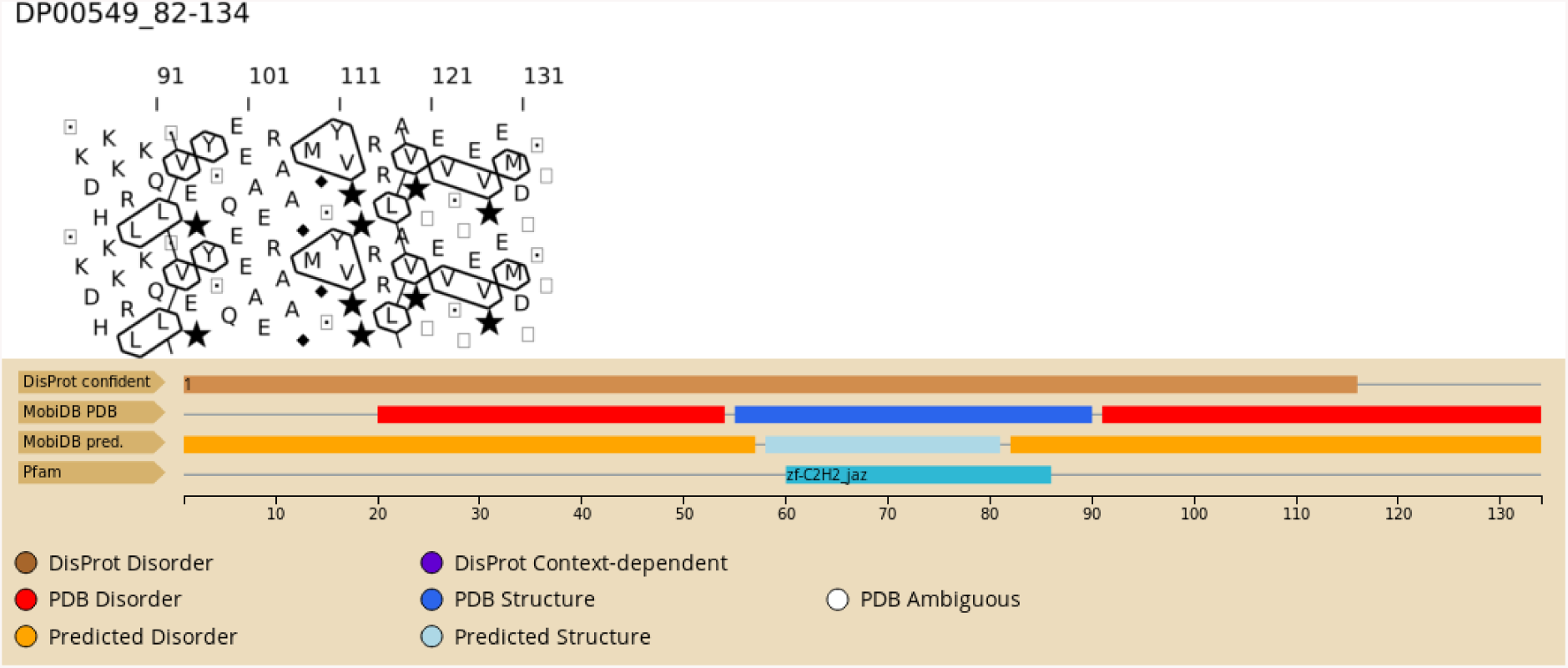
Disordered regions with high HCA score - example n°2. The HCA pattern displayed by this disordered region is similar to pattern of foldable regions. Clusters of several hydrophobic residues can be seen close together. The disordered region, (82-134) corresponds to the C-terminal of human Zinc finger protein 593 (UniProt O00488).

### Examples of PDB sequences with high HCA scores

Figures S7 and S8 show two examples of HCA plots for sequences extracted from the PDB. Figure S7 corresponds to the HCA plot of the Archeoglobus fulgidus VapC ribonuclease (Uniprot O28590, amino acids 1 to 156) whose 3D structure has been solved X-ray crystallography (PDB entry 1W8I). This ribonuclease is involved in a toxin-antitoxin module with toxin activity [1] and includes one known domain (amino acids 3 to 127 corresponds to the PFAM domain PIN, PF01850). The corresponding structure includes 9 long *α*-helices with 40% of hydrophobic amino acids. The dense network of HCA clusters visible in Fig. S7 is typical of globular proteins and the long *α*-helices can be visualized as horizontal clusters on the 2D HCA plot.

**Figure S7:**
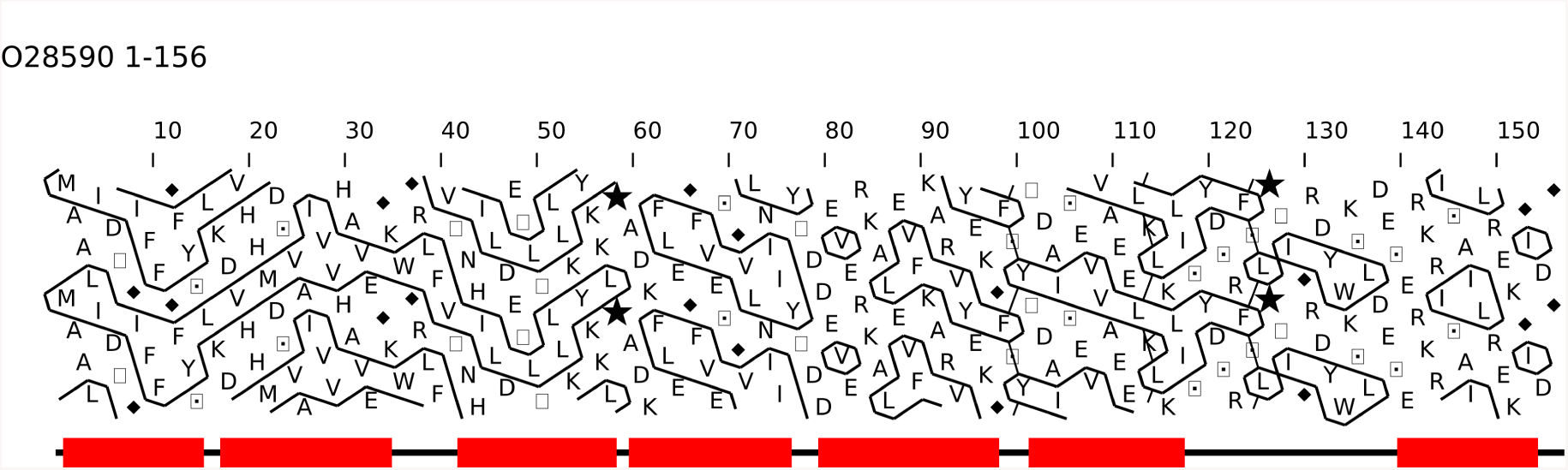
PDB sequence with an high HCA score - example n°1. This sequence (Uniprot 028590, PDB 1W8I), including amino acids 1 to 156, corresponds a typical globular protein. (*α*-helix: red rectangle, annotations extract from the experimental 3D structure 1W8I).

Figure S8 is another example of a globular protein HCA plot, i.e. with a high HCA score. The figure shows the HCA plot of the mature mouse interferon beta (Uniprot P01575, amino acids 22 to 181, PDB entry 1WU3 [26]). The protein is made of one domain (Pfam amino acids 27 to 179 (PF00143)), including 5 long *α*-helices with 42% of hydrophobic residues. As for Fig. S7, the protein contains large hydrophobic clusters, typical of regular secondary structures, separated by loops.

**Figure S8:**
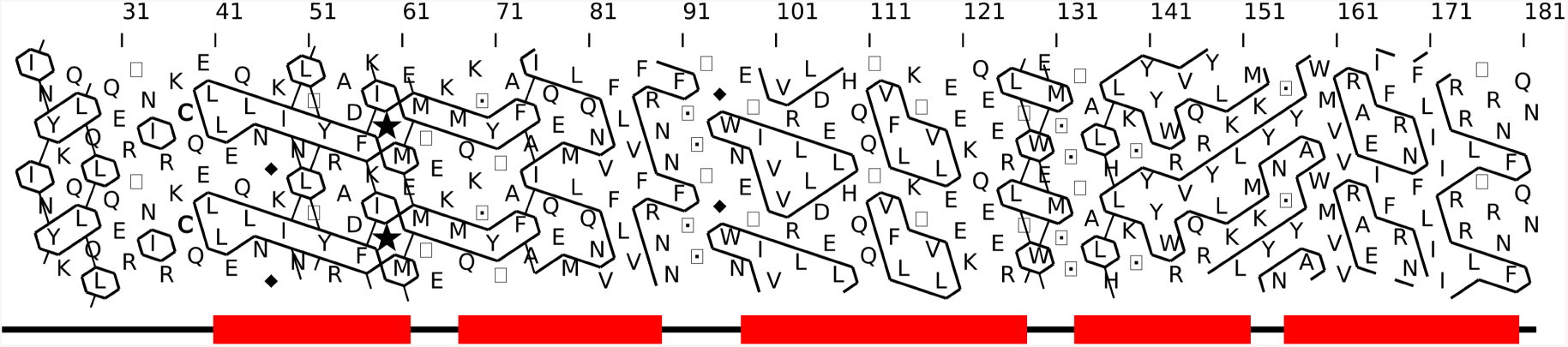
PDB sequence with an high HCA score - example n°2. The sequence region (Uniprot P01575, PDB entry 1WU3), including amino acids 22 to 181, corresponds to the PDB structure 1WU3 a typical globular protein. (*α*-helix: red rectangle, annotations extracted from the experimental 3D structure 1WU3).

### Examples of PDB sequences with low HCA scores

Figures S9 to S10 are two examples of PDB sequence HCA plots with low HCA scores. Fig. S9 corresponds to the N-terminal domain (amino acids 50 to 174) of the nucleoprotein from human SARS coronavirus (Uniprot protein P59595, PDB entry 2OFZ [16]). This nucleoprotein has RNA binding activity, packaging the positive strand of the human SARS coronavirus RNA genome into a helical ribonucleocapsid [27]. The RNA binding activity is mediated by the region encompassing amino acids 45 to 181, such binding activity is usually mediated by a high level of flexibility. The full-length protein is made of one or two protein domains according to the Pfam database (PF00937, from 15 to 378) or the SUPERFAMILY database (SSF110304, from 28 to 181 and SSF103068, from 252 to 365). The 3D structure of the first SSF domain is made of four *β*-strands (from amino acids 61 to 59, 84 to 91, 102 to 113, and 130 to 135) and one small *α*-helix (50 to 57), with a large number of flexible loops around the *β*-sheet core (Saikatendu et al., 2007). According to the coverage of the sequence by large loops, this protein domain has a lower percentage of hydrophobic residues (26%), than the regularly admitted of 30% limit, characteristic of globular domains.

Fig. S10 is another example of PDB protein sequence with low HCA score. The HCA plot represents a sequence segment (from amino acids 500 to 629) of the the Staphylococcus aureaus surface protein G

**Figure S9:**
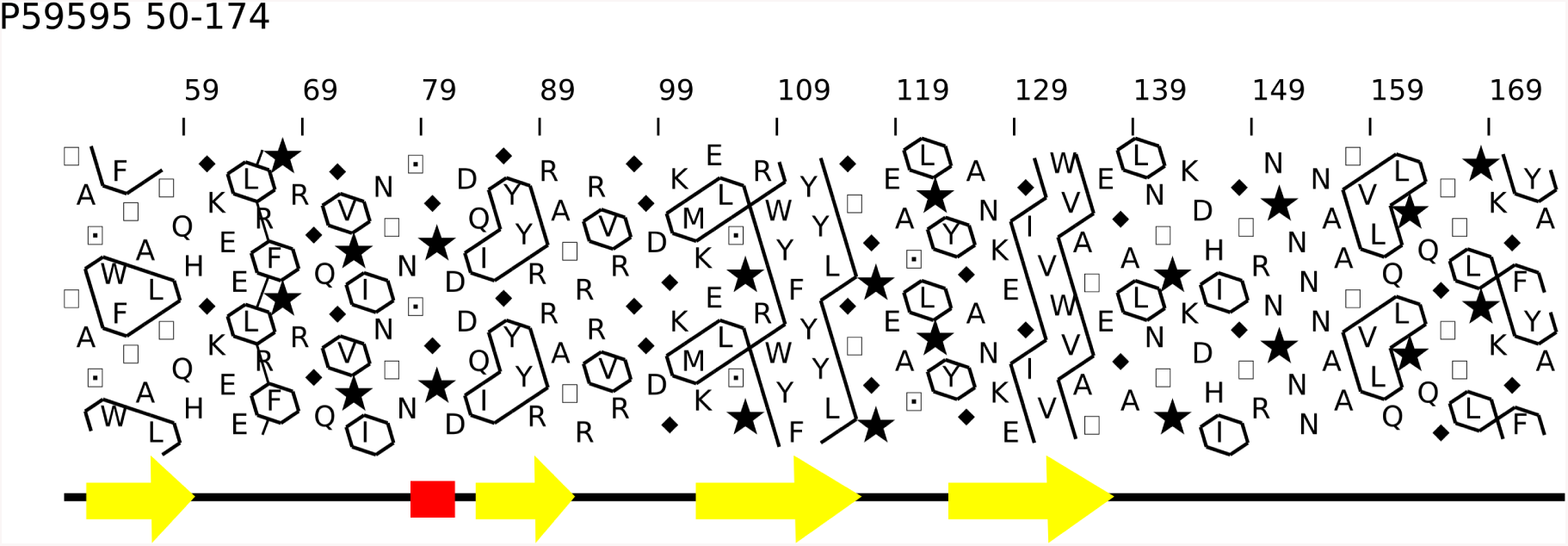
PDB sequence with a low HCA score - example n°1. The sequence region (Uniprot P5995), amino acids 49 to 174, PDB structure 2OFZ), corresponds to a RNA-binding domain, having large, flexible loops and a few regular secondary structures, constituting a *β*-sheet core. (*α*-helix: red rectangle *β*-strands: yellow arrows, annotation extracted from the experimental 3D structure 2OFZ).

 (SasG) (Uniprot Q2G2B2 sequence, PDB entry 5DBL). The full-length protein is made of 19 domains. The sequence starts with a signal peptide motif, followed by pairs of G5 domain/E domain (Pfam PF04650, PF17041) and ends by a cell wall anchor domain (PF00746). Amino acids 501 to 548 and 547 to 629 corresponds to a pair of E domain/G5 domain. The G5 domain has only a few conserved amino acids and is supposed to have an adhesive function [2]. As assessed by the presence of small clusters and a relatively weak percentage in strong hydrophobic amino acids, approximatively one half of the SasG repetitive regions are intrinsically unfolded in isolation, but fold in the context of neighboring folded G5 domains, highlighting the role of the intrinsically disordered region of the E/G5 domain pair as a key factor for the cooperative folding multidomain protein [14]. Once folded, the two domains form an elongated structure, made of small beta strands which correspond on the HCA plot to small clusters. The small *β*-strands form triplets-stranded *β*-sheets connected by collagen-like triple helical regions. In this particular case, several threonine are found included in *β*-beta strand.

**Figure S10:**
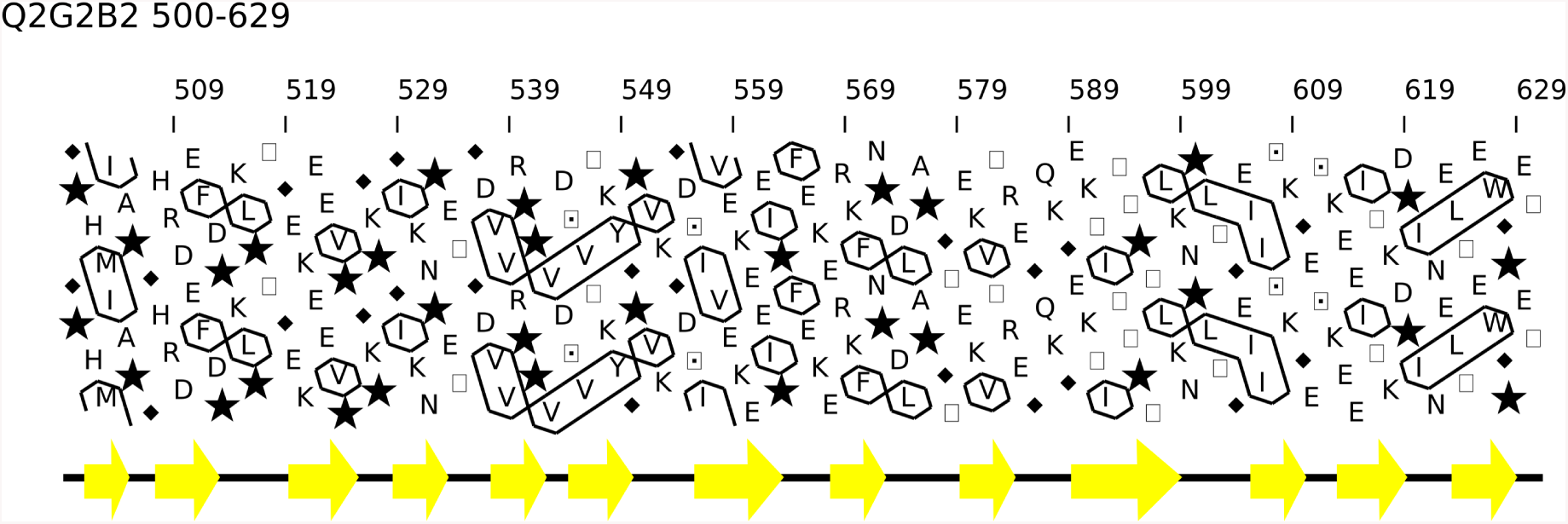
PDB sequence with low HCA score - example n°2. The sequence region, (Uniprot Q2G2B2, amino acids 500 to 629, PDB structure entry 5DBL), corresponds to a pair of E/G5 domains of the S. aureus surface protein G. (*β*-strand: yellow arrow, annotations extracted from the experimental 3D structure 5DBL).

